# A Mass Spectrometry-Based Interactome Study on Anisomelic Acid and Its Effects on HPV16 E7

**DOI:** 10.1101/2025.11.14.688426

**Authors:** Michael Santos Silva, Leila S. Coelho-Rato, Navid Delshad, Tatiana Tarkhova, Joakim Edman, Preethy Paul, Annika Meinander, John E. Eriksson

**Author notes:** Corresponding author at: Faculty of Science and Engineering, Cell Biology, Åbo Akademi University, 20520 Turku, Finland., e-mail addresses (M. Santos Silva), (J. E. Eriksson).

## Abstract

Human papillomavirus (HPV)-driven epithelial cancers rely on actions of viral oncoproteins, notably E7, which disrupts cell cycle control through inactivation of the tumor suppressor pRb. While prophylactic vaccines prevent infection, effective therapeutics for existing HPV-associated malignancies remain lacking. Here, we investigate the natural diterpenoid anisomelic acid (AA) and its mechanism of promoting HPV16 E7 degradation. Using cellular thermal shift assay (CETSA), we demonstrate that AA binds to E7, increasing its thermal stability and suggesting a direct molecular interaction. Mass spectrometry-based interactome analysis revealed that AA treatment reshapes the E7 interactome, enriching for E3 ubiquitin ligases (UBR4, TRIP12, TRIM28), the scaffold protein SQSTM1, and HSP40/HSP70 chaperone complexes involved in protein quality control and misfolded protein signaling. These data support a model in which AA binding induces conformational changes in E7 that enhance its recognition by ubiquitin-proteasome and chaperone-mediated degradation pathways. Our findings provide mechanistic insight into AA’s antiviral mode of action and highlight its potential as a lead compound for targeted degradation of HPV oncoproteins.

## 1 Introduction

Human papillomaviruses (HPVs) are small double-stranded DNA viruses capable of infecting epithelial tissues(1,2). High-risk HPV types are recognized as a major etiological factor in the development of cervical cancer and other epithelial malignancies(2). Cervical cancer, caused by HPV, continues to rank among the most common cancers affecting women worldwide(3). Although HPV is often associated with cervical carcinomas and women, high-risk HPV types also contribute significantly to the development of anogenital, anal, and oropharyngeal cancers, which can affect men as well(4).

Despite extensive vaccination efforts and significant advances in early detection, HPV-driven malignancies remain a major global health challenge. The currently available prophylactic vaccines efficiently prevent new HPV infections, but they are ineffective in individuals with already established infections or malignant transformation. Thus, there remains an unmet need for therapeutic strategies that selectively target HPV-infected cells and viral oncoproteins without affecting normal tissues.

High-risk HPVs rely primarily on two viral proteins, E6 and E7, to promote malignant transformation. These oncoproteins modulate different essential signaling mechanisms in infected cells(5). The E6 protein is mainly known for facilitating the degradation of the tumor suppressor p53 by associating with the E6-associated protein (E6AP/UBE3A)(6,7). In parallel, E7 primarily targets the retinoblastoma protein (pRb), a key regulator of the cell cycle(8). This interaction disrupts cell cycle checkpoints, freeing E2F transcription factors to drive uncontrolled cell proliferation, a hallmark of HPV-driven oncogenesis(9). Besides its well-characterized role in pRb degradation, the E7 oncoprotein also contributes to genome instability, metabolic reprogramming, and immune evasion through its impact on multiple cellular pathways, underscoring its central role in HPV-mediated carcinogenesis(10). Given these essential and virus-specific functions, both E6 and E7 are considered highly attractive therapeutic targets.

Previous studies from our group identified anisomelic acid (AA), a natural compound isolated from *Anisomeles malabarica*, as a promising small-molecule candidate capable of downregulating HPV16 E6 and E7 proteins(11). Treatment with AA restored p53 and p21 levels and induced cell cycle arrest and apoptosis selectively in HPV-positive cells, while exhibiting minimal cytotoxicity in HPV-negative cells. These findings demonstrated that AA selectively targets HPV-transformed cells and exerts its effects through a viral protein-dependent mechanism. Moreover, inhibition of the proteasome with MG132 prevented the AA-mediated reduction in E6, indicating that AA promotes the degradation of HPV oncoproteins through a proteasome-dependent pathway(11). The subsequent mass spectrometry-based study from our group revealed that AA induces E6 degradation via recruitment of specific E3 ligases, providing direct evidence that AA acts through the ubiquitin–proteasome system(12). While the mechanism underlying E6 degradation has thus been elucidated, the molecular details of how AA affects the E7 oncoprotein remain unexplored. Considering the interdependence of E6 and E7 in maintaining HPV oncogenic potential, it is plausible that AA might also modulate E7 stability and promote its degradation through similar or converging pathways. To investigate this possibility, we employed a mass spectrometry-based proteomics approach to map the E7 interactome under AA treatment and identify the host proteins mediating its degradation. In parallel, we used cellular thermal shift assays (CETSA) to determine whether AA binds to E7, potentially inducing conformational changes that enhance its susceptibility to ubiquitination. Through these complementary approaches, we aimed to elucidate how AA influences E7’s stability, identify the cellular machinery involved in its turnover, and uncover the broader molecular mechanisms by which AA exerts its antiviral and antitumor effects.

Here, we show that AA binds to E7 and enhances its ubiquitination, leading to its proteasomal degradation. Using mass spectrometry-based interactome profiling, identified we identify key E3 ubiquitin ligases (UBR4, TRIP12, and TRIM28), the scaffold protein SQSTM1, and HSP40/HSP70 family members as potential mediators of E7 degradation. Together, these findings provide new insight into how anisomelic acid modulates the host cell’s protein quality control machinery to specifically target HPV oncoproteins for degradation. This work extends our understanding of AA’s mode of action and establishes a foundation for developing targeted therapeutic strategies for HPV-associated malignancies.

## 2 Experimental Procedures

### 2.1 Cell Culture, Treatments, and Transfections

SiHa (HTB-35, ATCC) were grown in standard conditions in Dulbecco’s Modified Eagle Medium (DMEM). The medium was supplemented with 10% fetal bovine serum (FBS; S1810, Biowest), penicillin (100 U/mL), streptomycin (100 μg/mL, P0781, Sigma), and L-glutamine (2 mM, X0550, Biowest).

For transient transfection assays, SiHa cells were electroporated with plasmids encoding N-CMV-HA-FLIP-16E7 (a kind gift from Prof. Karl Munger), or pCMV-Neo-Bam (16440, Addgene) as a control. For each transfection, a 0.4 cm Gene Pulser/MicroPulser Electroporation Cuvette (1652081, Bio-Rad) was used, containing 3 x 10^6^ cells in 800µL Opti-MEM (31-985-062, Gibco) and 10 µg of plasmid. Cells were electroporated using a single electric pulse (220V, 975 µF) from a Gene Pulser II electroporator (Bio-Rad) and subsequently cultured in DMEM. 24 hours after transfection, cells were exposed to either anisomelic acid (40 µM) or DMSO as a control, with or without a 2-hour pre-treatment with the proteasome inhibitor MG132.

Stable normal oral keratinocytes (NOK) cell lines expressing HPV16 oncogenes (NOK16) and their parental negative control (pWPI) were kindly gifted by Dr. Martina Niebler (DKFZ, Germany). Cells were cultured according to the reported protocol(13). To assess the optimal AA concentration, NOKs were treated with 10, 20, or 40 µM of AA for 24 hours, and samples were analyzed by western blot.

### 2.2 SDS-PAGE and Western Blotting

Cells were harvested and lysed in 1x Laemmli sample buffer (#1610737, Bio-Rad) supplemented with 2.5% β-mercaptoethanol. Lysates were heated at 98°C for 5 minutes and were separated on 4–20% Mini-PROTEAN® TGX precast gels (#4561093 and #4561096, Bio-Rad) using 1x Tris/Glycine Buffer (#1610771, Bio-Rad) containing 0.1% SDS. Electrophoresis was performed for 40 minutes at 200 V. Following electrophoresis, proteins were transferred onto methanol-activated polyvinylidene difluoride (PVDF) membranes (#10600023, Amersham) with a wet transfer system containing 1x Tris/Glycine Buffer with 20% methanol. Wet transfer was performed at 400 mA for 1 hour. Membranes were blocked for 1 hour at room temperature in 5% milk prepared in TBS with 0.3% Tween-20, washed, and subsequently incubated overnight at 4°C with primary antibodies. After washing for 20 minutes with TBS or PBS containing 0.3% Tween-20, membranes were incubated for 1 hour at room temperature with secondary antibodies. Protein detection was performed by chemiluminescence (Amersham RPN2109 and RPN2236 mixed at a 2:1 ratio), and membranes were imaged using the iBright Imaging System (CL1000, Thermo Fisher Scientific). The following primary antibodies were diluted 1:1000 in TBS or PBS 0.3% Tween20 3% BSA 0.02% sodium azide: HA (3724S, CST), HSC70 (ADI-SPA-815, Enzo), Strep-tag II (ab76949, Abcam), pRb (9309, CST) and β-actin (5125S, CST),. The secondary antibodies used for this study were diluted 1:10000 in TBS or PBS 0.3% Tween20 5% milk and included: anti-rabbit (W401B, Promega), anti-mouse (W4021, Promega), and anti-rat (NA935, GE).

### 2.3 Tandem Ubiquitin Binding Entities (TUBEs) pulldown

Post-transfection, SiHa cells were treated with MG132 and anisomelic acid (AA), as described previously. Cells were harvested by trypsinization and lysed on ice for 10 minutes using a lysis buffer (20 mM disodium phosphate, 20 mM monosodium phosphate, 1% NP-40, 2 mM EDTA, 1% SDS, 1 mM DTT, 5 mM N-ethylmaleimide, 5 mM chloroacetamide, and 1x protease and phosphatase inhibitors [A32961, Thermo Fisher Scientific]). Lysates were diluted to a final SDS concentration of 0.1% and were centrifuged for 10 minutes at 12,000 rpm at 4°C. 6 µL of TUBE reagent and 20 µL of a 50% glutathione bead slurry were added to the supernatant. The mixtures were incubated at 4°C for 2 hours with gentle rotation, followed by four washes using PBS containing 0.1% Tween-20. After washing, beads were boiled in 50 µL of Laemmli sample buffer and analyzed by SDS-PAGE and western blotting.

### 2.4 Co-immunoprecipitation

For mass spectrometry analysis, NOK cells were used for co-IP. For each treatment condition (untreated, AA, AAMG), four biological replicates were processed. Following treatment, cells were washed with PBS and lysed in 500 µL of lysis buffer (50 mM Tris-HCl pH 7.5, 150 mM NaCl, 0.5% IGEPAL CA-630, supplemented with 1x protease and phosphatase inhibitors). Lysis was performed on rotation for one hour at 4°C. Lysates were centrifuged at 13,000 g for 10 minutes at 4°C. Ten percent of each supernatant was reserved as input control, and the remaining lysate was used for immunoprecipitation. Dynabeads™ MyOne™ Streptavidin C1 (50 µL per sample, 65001, Invitrogen^TM^) were washed three times with PBS and added to the cleared lysates. Samples were incubated overnight at 4°C on rotation. After incubation, beads were washed three times with PBS. Samples were boiled in 50 µL of Laemmli for western blot or processed further for mass spectrometry.

### 2.5 Cellular Thermal Shift Assay (CETSA)

15 cm dishes were used to culture NOK16 cells overnight, in experimental medium consisting of 22.5% DMEM, 72.5% F-12, 5% FBS, 5 µg/mL insulin, penicillin/streptomycin, 10 ng/mL epidermal growth factor, 24 µg/mL adenine, 8.4 ng/mL cholera toxin and 0.4 µg/mL hydrocortisone. The following day, cells were treated for one hour with either DMSO or 20 µM AA. After treatment, cells were washed with PBS, detached by trypsinization, centrifuged at 300 g for 3 minutes, washed with PBS, and resuspended in 1 mL of PBS. Aliquots of 100 µL were subjected to heat treatment at different temperatures (43, 47, 50, 53, 57, and 60°C) for 3 minutes using a thermal cycler, followed by a 3-minute incubation at room temperature. Dry ice was used to snap-freeze the sample, which were stored at −80°C until further processing. For western blot analysis, cell lysis was performed by three consecutive freeze-thaw cycles on dry ice and vortexing between each cycle. Lysates were centrifuged for 20 minutes at 20,000 g at 4°C, and the resulting supernatants were boiled in 2x Laemmli buffer.

### 2.6 Mass Spectrometry

Lysis buffer (50 mM Tris-HCl, pH 7.5, 150 mM NaCl, 0.5% IGEPAL CA-630, supplemented with 1x protease and phosphatase inhibitors), ice-cold PBS and MilliQ water were used three times each to wash co-IP samples. Subsequently, proteins bound to the beads were digested at 27°C for 30 minutes on a thermomixer set to 800 rpm in a digestion buffer containing 5 µg/mL trypsin in 6 volumes of 2 M urea, 25 mM ammonium bicarbonate (ABC) pH 7.5. After digestion, supernatants were collected into clean tubes, while the beads were washed twice with 2.5 volumes of 2 M urea, 25 mM ABC, pH 7.5, containing 1 mM DTT. Wash fractions and the initial digests were pooled and further digested overnight at room temperature. The following day, 20 µL of 5 mg/mL iodoacetamide (IAA) was added to the samples and incubated for 30 minutes in the dark. Trifluoroacetic acid (TFA) was used to acidify samples before they were desalted using a Sep-Pak C18 96-well plate (40 mg, 186003966, Waters) according to the manufacturer’s guidelines. Samples were dried in a speed-vacuum centrifuge and stored at −80°C until further use. Prior to LC-MS/MS analysis, 0.1% formic acid was used to dissolve the peptides. For each run, 400 ng of peptide material was injected.

Peptides were analyzed using an EASY-nLC 1200 HPLC system (Thermo Fisher Scientific) coupled to an Orbitrap Fusion Lumos mass spectrometer (Thermo Fisher Scientific) equipped with a nano-electrospray ionization source. Samples were first loaded onto a trapping column (100 µm internal diameter × 2 cm) and later separated on an analytical column (75 µm internal diameter × 15 cm), both packed in-house with ReproSil-Pur C18-AQ resin (3 µm, 120 Å; Dr. Maisch HPLC GmbH). Peptides were eluted using a 60-minute gradient as follows: 8–21% solvent B over 28 minutes, 21–36% B over 22 minutes, 36–100% B over 5 minutes, and a final wash at 100% B for 5 minutes. Solvent B consisted of 80% acetonitrile and 20% water with 0.1% formic acid, delivered at a constant 300 nL/min flow. Full MS scans were acquired in the Orbitrap across 300–1750 m/z range at a resolution of 120,000. The automatic gain control (AGC) target was set to 700,000 with a maximum injection time of 50 ms. Precursors with charges greater than +2 and intensities above 25,000 were selected for fragmentation within a 2.5-second cycle. These ions were isolated using a 1.6 Da window in the quadrupole and fragmented using higher-energy collisional dissociation (HCD) at 27% normalized collision energy. MS/MS spectra were collected in the Orbitrap at a resolution of 30,000, with an AGC target of 50,000 and maximum injection time of 75 ms. Dynamic exclusion was set to 35 seconds with a 10 ppm mass tolerance.

Raw LC-MS/MS files were processed using MaxQuant (version 2.0.1.0)(14,15) or FragPipe (version 21.1) with associated tools: msFragger (4.0), IonQuant (1.10.12), Philosopher (5.1.0), and Python (3.9.13)(16–18).

For MaxQuant analysis, searches were performed against the UniProt HPV proteome (UP000009251, 9 entries) and the human reference proteome (UP000005640, 78,120 entries), both downloaded on August 1, 2021, including MaxQuant contaminants. Search parameters included methionine oxidation and N-terminal acetylation as variable modifications, carbamidomethylation as a fixed modification, and trypsin digestion allowing up to two missed cleavages. Mass tolerances were set at 20 ppm for the first search and 4.5 ppm for the main search. Label-free quantification (LFQ) was enabled with a minimum ratio count of 2, classic normalization, and quantification based on unique + razor peptides. The false discovery rate (FDR) was set at 1%, with match between runs enabled (match time window: 2 minutes, alignment time window: 20 minutes). Downstream analysis was carried out using Perseus (version 1.6.15.0). Reverse sequences, contaminants, and proteins identified only by site were removed. Data were transformed and filtered to retain proteins quantified in at least three replicates per group. Venn diagrams were created using InteractiVenn(19), applying the median of replicate groups from different analysis stages. Data imputation followed a normal distribution (width 0.3, downshift 1.8) across the full matrix and was filtered for ANOVA significance (p < 0.05). Z-score normalized data with Euclidean distance was used for hierarchical clustering heatmaps. Ontology analysis was done using ShinyGO 0.80(20) with a human background, reporting the top 10 enriched biological processes with an FDR < 0.05 and a minimum pathway size of 2. Perseus was used to create the Hawaii plot using Pearson correlation. Non-imputed datasets were filtered using a ratio of E7/pWPI > 1.05.

For FragPipe searches, the UniProt human database (UP000005640) was downloaded on September 5, 2023, containing 40,912 entries with 50% reverse decoy sequences. The default LFQ-MBR workflow was applied with the following parameters: precursor and fragment mass tolerances at 20 ppm, trypsin specificity allowing up to two missed cleavages, methionine oxidation as a variable modification, and cysteine carbamidomethylation as fixed. FDR thresholds were set at 1% for both protein and peptide levels. Downstream data analysis was conducted in Perseus using the same approach as for MaxQuant. Output files (reprint.int) were processed through CRAPome(21) for SAINT(22) analysis, using default user control settings.

## 3 Results and Discussion

### 3.1 Anisomelic acid promotes ubiquitination and degradation of HPV16 E7

To assess how AA affects E7 protein levels, we used SiHa cells transiently transfected with FLAG-tagged HPV16 E7. After 24 hours, cells were treated with AA in the presence or absence of the proteasome inhibitor MG132. Western blot analysis showed a clear decrease in E7 protein levels following AA treatment, an effect prevented by the presence of MG132 co-treatment (Figure 1A). These results indicate that AA promotes proteasome-dependent degradation of E7, consistent with our previous observations for E6(11,12).

**Figure 1.**
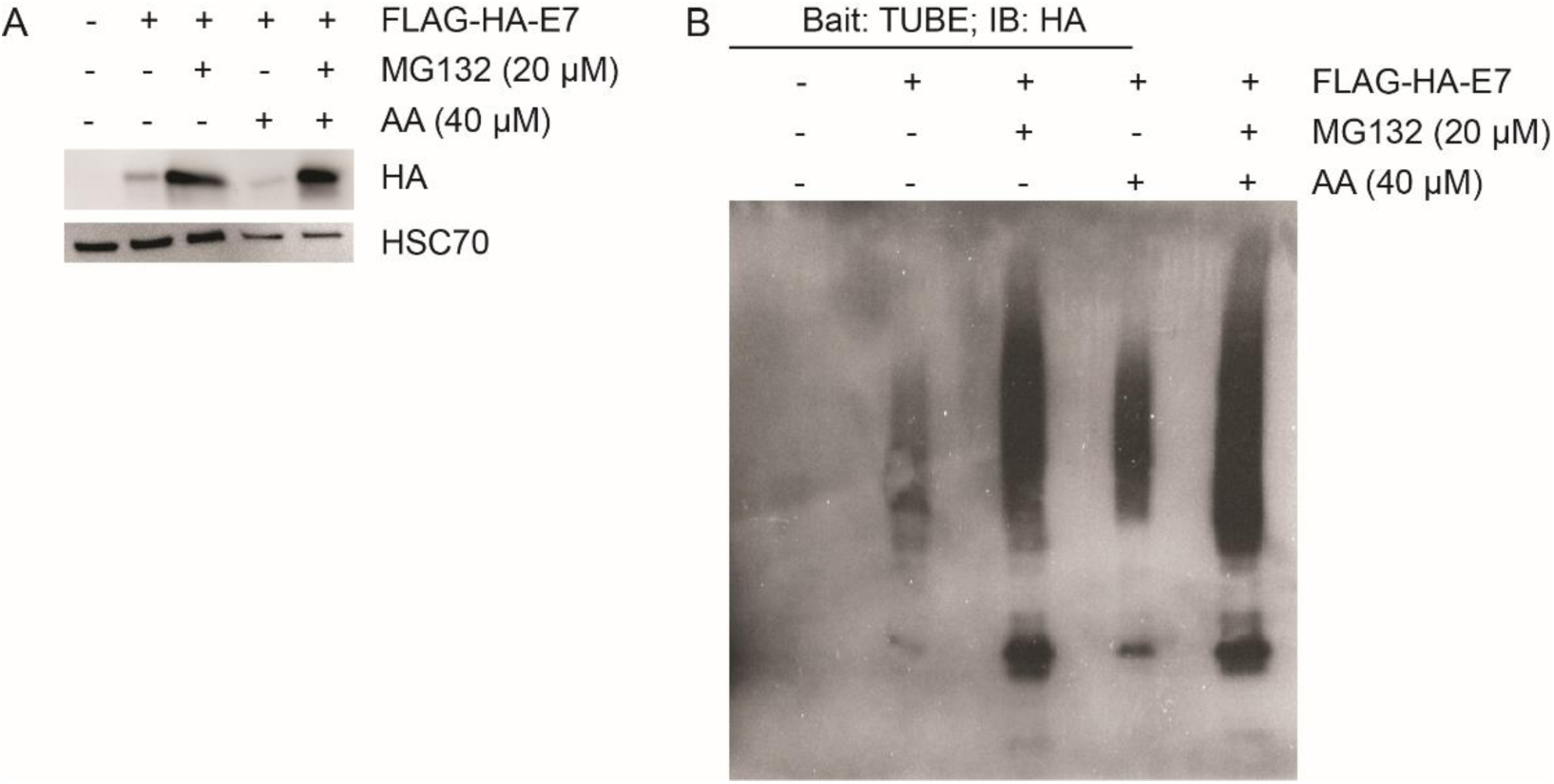
HPV16 E7 protein is ubiquitinated and degraded in the presence of AA. a) Western blot was performed on transfected SiHa cells with FLAG-HA-tagged HPV16 E7. 24 hours after transfection, cells were treated with MG132 for 2 h and/or with AA for 24 h. HSC70 served as the loading control. b) TUBE pulldown was conducted on transfected SiHa cells after treatment with MG132 (2 h) and/or AA (24 h).

Given the central role of ubiquitination in targeting proteins for proteasomal degradation, we next assessed whether AA influences the ubiquitination of E7. Using the same transfection and treatment protocol, we performed tandem ubiquitin-binding entity (TUBE) pulldown assays to enrich ubiquitinated proteins. The results demonstrated increased ubiquitination of E7 in AA-treated samples compared to untreated controls, while co-treatment with MG132 led to further accumulation of ubiquitinated E7 (Figure 1B). These findings confirm that AA enhances E7 ubiquitination and degradation via the ubiquitin-proteasome pathway. The involvement of Cullin 1– and Skp2-containing E3 ligases in E7 ubiquitination and degradation has been reported previously(23–25). However, these studies focused on steady-state conditions, suggesting that maintaining a balance in E7 protein levels is crucial for HPV infection. Interestingly, the ectopic expression of suppressor of cytokine signaling-1 (SOCS1), a JAK tyrosine kinase binding protein, has been shown to promote E7 degradation by direct binding and enhancing its ubiquitination in SOCS-box-dependent manner(26). Furthermore, activation of STING-TBK1 (Stimulator of Interferon Genes and TANK-binding kinase 1), key players in innate immune response against viral infections, has also been reported to promote ubiquitination and degradation of E7 oncoprotein(27). Lastly, docosahexaenoic acid was shown to induce E7 downregulation by increased reactive oxygen species (ROS) production(28), although the precise mechanism was not further investigated.

Considering these examples, we hypothesized that AA could either modulate upstream regulators, such as E3 ligases, or directly bind E7, altering its conformation and rendering it more susceptible to ubiquitination. To investigate whether AA directly interacts with E7, we performed a cellular thermal shift assay (CETSA)(29) using NOK cells stably expressing HPV16 STREP-tagged E7 (NOK16)(13). The CETSA method is based on the principle that direct ligand binding stabilizes the target protein against thermal denaturation. Our experiment revealed that E7 exhibited increased stability in the presence of AA, as indicated by stronger bands at 47 °C, 50 °C and 53 °C compared to DMSO controls (Figure 2A). These results suggest a direct interaction between AA and E7, potentially inducing conformational changes that facilitate E7 ubiquitination and degradation. Further studies, such as surface plasmon resonance or cryo-EM, could help define binding kinetics and structural interfaces, but the intrinsic instability of recombinant E7 has thus far limited these analyses.

**Figure 2.**
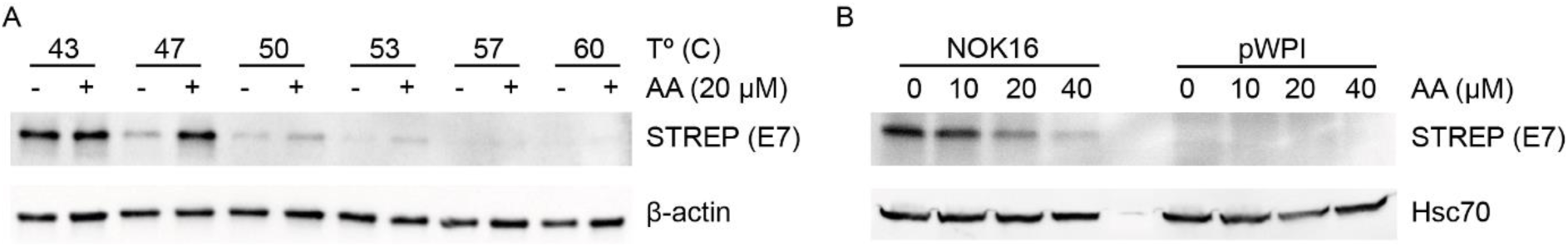
AA downregulates E7 in NOKs stably expressing STREP-tagged HPV16 E7. a) CETSA was preformed by trypsinizing and harvesting NOK cells, followed by heating at temperatures ranging from 43-60°C in the presence or absence of AA. Western blot was used to assess E7 levels, using β-actin as a loading control. b) Western blot analysis was performed on NOK16 and pWPI treated for 24 h with different concentrations of AA. HSC70 was used as a loading control.

To gain further insight into the degradation pathway and identify potential candidate E3 ligases involved in E7 ubiquitination, we employed mass spectrometry-based proteomics using NOK cells stably expressing HPV16 STREP-tagged E7. Before proceeding with the interactome analysis, we assessed the impact of AA on E7 protein levels in this system. Western blot analysis showed that E7 levels decreased in an AA-dose dependent matter (Figure 2B), supporting the use of 20 µM AA for subsequent proteomic experiments.

Our findings indicate that AA facilitates the degradation of HPV16 E7 by enhancing its ubiquitination and subsequent targeting to the proteasome, consistent with previous reports describing the impact of AA on endogenous E7 protein levels(13). Furthermore, the observation that AA produces similar effects in HPV-positive cervical cancer cells and in transformed oral keratinocytes suggests a broader mechanism of action across different HPV-positive cell types. This validates the use of NOK cells as a relevant model to investigate AA’s mechanism of action. Importantly, we demonstrate for the first time that AA can directly interact with E7, lending further support to the hypothesis that AA binding may increase E7’s susceptibility to proteasomal degradation.

### 3.2 Mass spectrometry analysis identifies E3 ligases and proteasomal subunits

To elucidate the molecular players involved in AA-mediated E7 degradation, we performed co-immunoprecipitation (co-IP) using STREP-beads, followed by LC-MS/MS analysis. This experimental strategy was adapted from our previous work(12). Briefly, four treatment conditions were analyzed (n=4): NOK pWPI (negative control), NOK16 untreated (control), NOK16 treated with AA (AA), and NOK16 treated with both AA and MG132 (AAMG).

To thoroughly evaluate the impact of AA on the E7 interactome and ensure high-quality data, we analyzed the mass spectrometry data using both MaxQuant and FragPipe pipelines. This dual analysis allowed us to cross-validate protein identifications and ensure consistency between database search algorithms. During data preprocessing, we excluded potential contaminants, reverse sequences, and proteins only identified by site. Proteins with a minimum of three valid values in at least one experimental group were kept and log₂-transformed for further analysis. This filtering resulted in identification of more than 450 proteins by MaxQuant and over 600 proteins by FragPipe (Figure S1A-B). AAMG-treated samples exhibited the highest number of identified interactors, as expected due to the inhibition of proteasomal degradation. The larger number of proteins recovered in the AAMG condition is consistent with stabilization of transient or low-abundance E7-associated complexes when degradation is blocked. Furthermore, FragPipe database search yielded a higher number of identified proteins, likely reflecting differences in search algorithms between the two platforms. To assess the effectiveness of the co-IP, we verified the detection of the retinoblastoma protein (pRb), a well-established E7 interactor, in our dataset. pRb was detected in E7-expressing samples, confirming that the co-IP successfully captured relevant interactors and that AA does not completely disrupt E7 interactions (Figure S1C-D). Interestingly, label-free quantification (LFQ) analysis showed that pRb levels were higher in AAMG and AA treatment conditions than in controls, consistent with pRb stabilization when E7 is degraded(30). While analyzing the overlapping proteins between treatment conditions, we found that most proteins were shared across conditions. In MaxQuant, we identified 16 proteins unique to AAMG-treated samples and 16 proteins shared between AA and AAMG (Figure S1E). In FragPipe, 1 protein was specific to AA, 27 were unique to AA+MG132, and 21 were shared between those conditions (Figure S1F). These results suggest that AA treatment remodels the E7 interactome.

Following initial data filtering, missing values were imputed using a normal distribution applied to the total matrix (width = 0.3, downshift = 1.8). Manual inspection of randomly selected rows confirmed imputation did not alter the data. Principal component analysis (PCA) showed clear clustering by treatment group (Figure 3A). The close grouping of biological replicates within each treatment group underscores the consistency and reproducibility of our experimental setup, while the distinct separation between treatments reflects treatment-specific alterations in the E7 interactome, particularly in response to AA and AA+MG132. To further explore protein enrichment among treatments, we performed hierarchical clustering of z-scored ANOVA-significant imputed data (Figure 3B). This analysis revealed three major protein clusters: proteins enriched in NOK16 samples, including those involved in translation, RNA splicing and processing, and ribonucleoprotein complex assembly; proteins enriched in AAMG-treated samples, associated with biological processes such as protein folding and translation; and proteins enriched in pWPI negative control samples, which served as a reference to identify and eliminate background proteins. These results offer mechanistic insight into how AA influences biological processes. Although we anticipated enrichment of proteasomal and ubiquitin-related processes, this was not evident with this clustering analysis. The enrichment of translation– and folding-related proteins may represent compensatory cellular stress responses triggered by E7 destabilization and loss of viral oncogenic control. Interestingly, our E6 study(12) showed proteasome-related enrichment, suggesting that AA-induced degradation of E6 may unbalance E6/E7 coordination, reactivating checkpoint and stress responses of the cell. This could potentially lead to E7 being targeted as a misfolded protein and thus being targeted for degradation.

**Figure 3.**
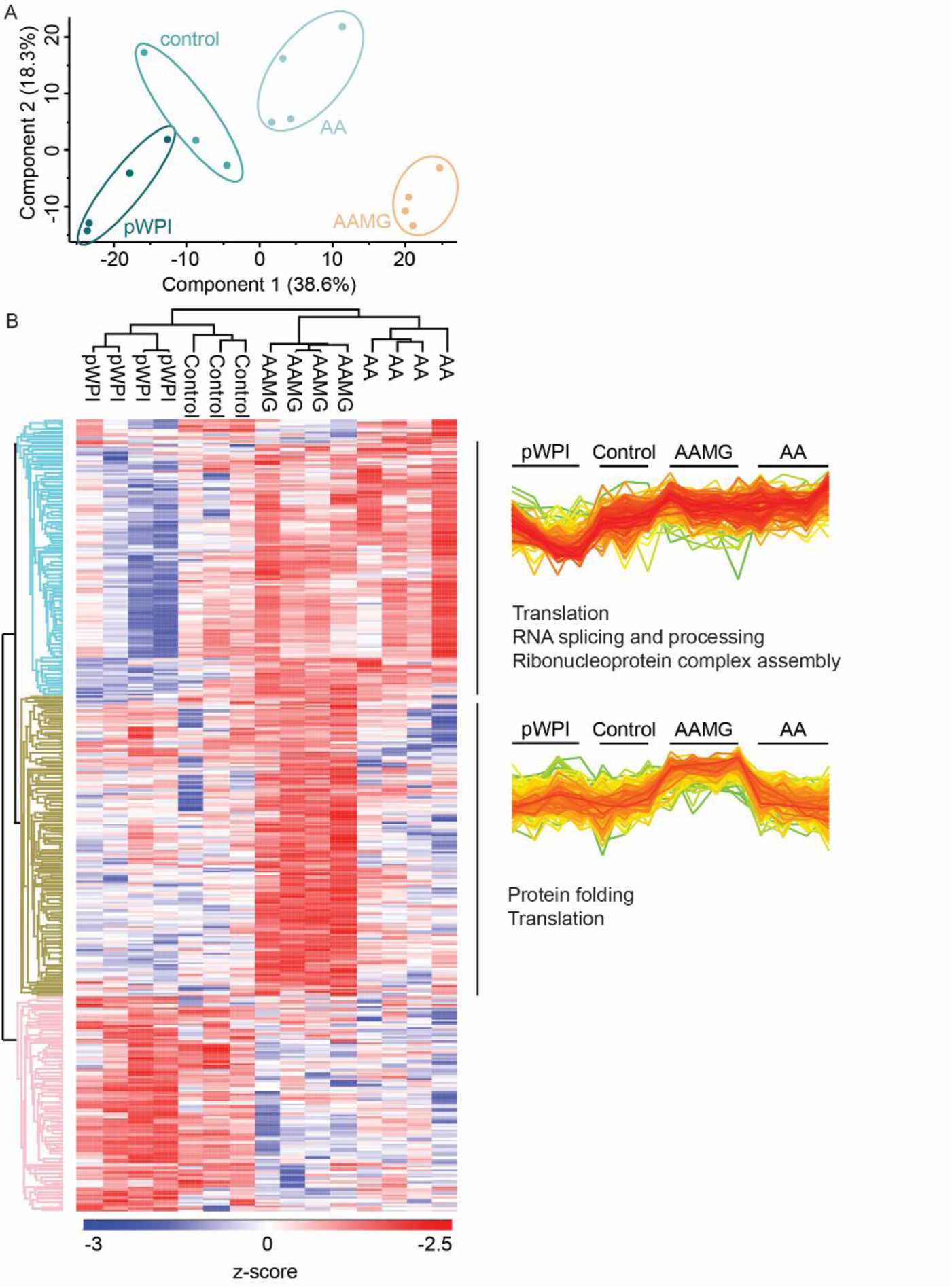
Imputed proteomics data reveal enrichment of translation and protein folding processes in AAMG-treated samples. a) Principal component analysis of all samples in this dataset. b) Hierarchical clustering was performed on imputed ANOVA significant z-scored data. Each row represents a protein, and each column corresponds to a sample. Functional enrichment of selected protein clusters was carried out based on Gene Ontology biological processes using ShinyGO 0.80.

Next, to further explore the dataset, volcano plots were generated for each condition, using a permutation-based FDR to identify Class A (high-confidence) and Class B (lower-confidence) interactors (Figure S2). No significant interactors were found in control samples. In contrast, both AA and AA+MG132 treatments yielded numerous Class A and B interactors. In AA and AAMG treatments, we identified several proteasomal subunits, including: PSMA3 and PSMC2 as Class B interactors in AA samples; PSMA3, PSMC1, PSMD2 and PSMD4 as Class A interactors, and PSMC2 as Class B interactor for AAMG samples. These findings support the hypothesis that AA promotes proteasome-dependent degradation of E7. In addition to proteasomal components, several E3 ubiquitin ligases also emerged as significant interactors. For instance, RBBP6 and UBR4 were both identified as Class A interactors for AA– and AAMG-treated samples. RBBP6 is known to bind to pRb, and to enhance cervical cancer malignancy by the JNK pathway(31,32). While E7-driven pRb degradation has been linked to calpain(33), RBBP6 could potentially mediate pRb degradation in AA-treated cells by targeting AA-bound E7. This may occur through conformational changes in E7 that increase its susceptibility to RBBP6-dependent ubiquitination. UBR4, known to associate with E7 and degrade PTPN14, contributes to the carcinogenic activity of HPV E7(34). Since UBR4 and E7 interact with each other, an AA-mediated conformational change of E7 could lead to UBR4 ubiquitinating E7 instead of PTPN14. Interestingly, UBR4 also appeared in our E6 dataset, suggesting it could regulate both oncoproteins. Additionally, TRIP12 was detected as a Class B interactor in the AAMG-treated group, while TRIM28 and NOSIP as Class A. Previous studies have shown that TRIP12 is downregulated in HPV-positive head and neck squamous cell carcinoma (HNSCC) and is regulated by p16, linking it to G1 arrest and impaired DNA repair(35,36). This raises the possibility that AA may affect the p16–TRIP12 axis, shifting TRIP12’s activity toward targeting viral proteins. Although no reports were found linking NOSIP to any cancer type, TRIM28 is upregulated in cervical cancers, contributing to cancer cell growth(37). Even though its interaction with HPV proteins remains unknown, this possibility cannot be ruled out yet, due to its apparent importance in cervical cancer cell growth(37). Overall, proteasome inhibition proved useful for stabilizing weak or transient interactions, enabling a more detailed characterization of the E7 interactome.

In conclusion, our mass spectrometry-based analysis demonstrates that AA treatment alters the E7 interactome by enhancing its association with proteins related to translation and protein folding. Co-treatment with MG132 increased the number of detected interactors, likely by preventing degradation of E7. Several candidate E3 ligases were identified, providing a foundation for future biochemical validation. Although the interactome analysis provided valuable insights, the large number of potential interactors could lead to challenges for downstream interpretation. To overcome this, we applied a filtering strategy aimed at refining the dataset to focus on the most biologically relevant proteins.

### 3.3 Data filtering reveals SQSTM1 and chaperones as potentially involved in E7 degradation

To reduce dataset complexity, we refined our analysis by excluding proteins that were more abundant in pWPI control cells compared to NOK16 cells. We applied a filter of NOK16/pWPI greater than 1.05 to eliminate likely background proteins, thereby generating a more focused subset for downstream analysis. As a quality control step, we cross-referenced our datasets with the CRAPome repository, which catalogs proteins frequently identified as nonspecific binders in affinity purification-mass spectrometry (AP-MS) experiments. The majority of proteins identified in our study were not listed as common contaminants, underscoring the specificity of our co-IP and mass spectrometry approach (Figure 4A and Figure S3A). To explore the functional implications of the filtered dataset, we performed gene ontology (GO) enrichment analysis. The top 10 enriched biological processes across treatment conditions are shown in Figure 4B (MaxQuant) and Figure S3B (FragPipe). Interestingly, AA and/or MG132 treatment did not substantially alter the overall categories of enriched biological processes, which remained dominated by translation, RNA processing, and protein folding pathways. However, while global biological processes remained stable in the presence of AA, individual protein changes may still occur.

**Figure 4.**
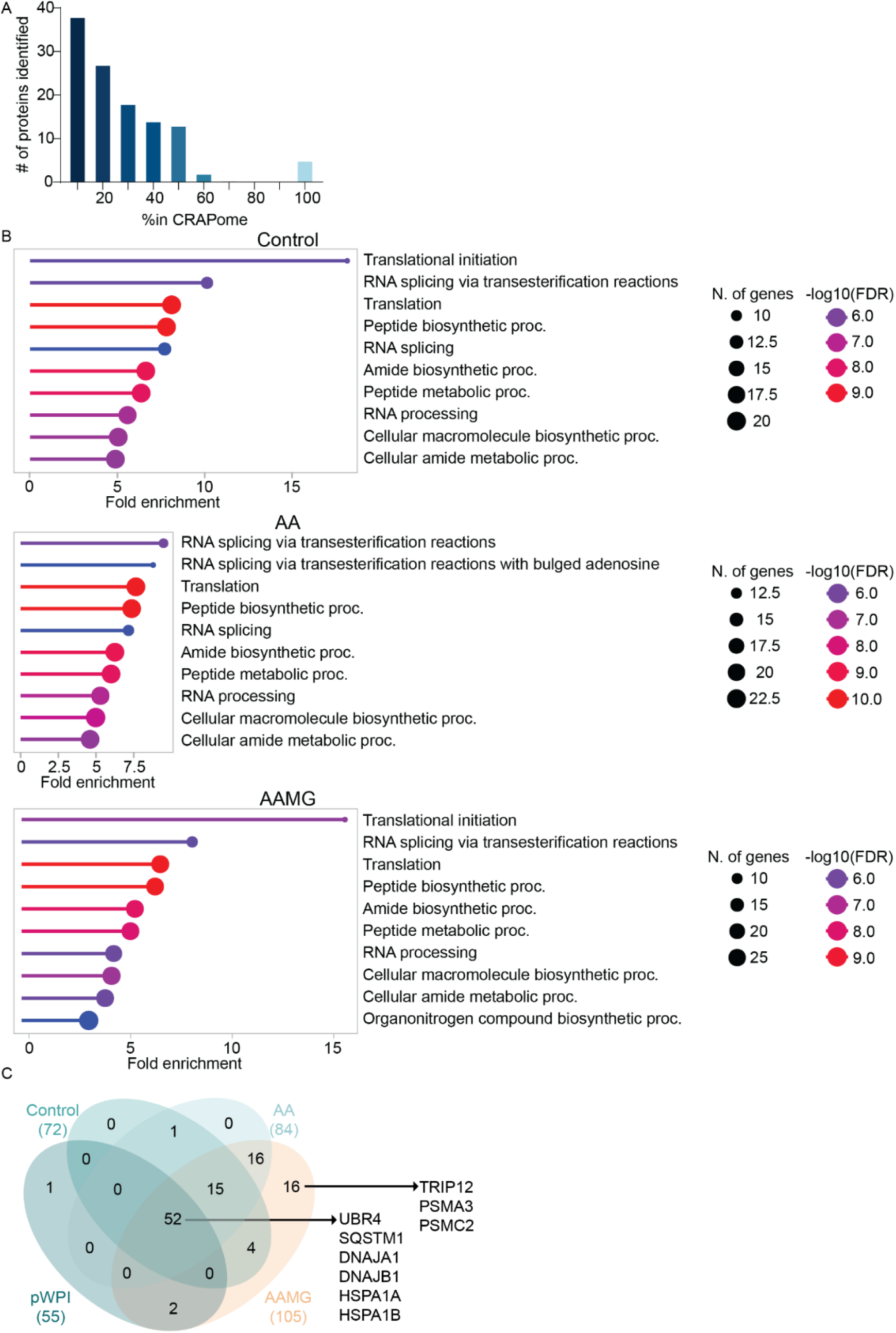
MaxQuant and Perseus analysis reveal enrichment of proteins from degradation and misfolded proteins pathways in NOK16. a) Bar plot showing how frequently proteins identified in this dataset appear in the CRAPome database. b) Dot plot summarizing GO biological processes across treatment conditions. Circle size indicates the number of proteins associated with each biological process, while color represents –log₁₀(FDR) values. c) Venn diagram comparing protein overlap between experimental groups, following exclusion of proteins with a NOK16/pWPI ratio below 1.05.

To assess this possibility, we compared the identified proteins across treatments using Venn diagrams. The MaxQuant-derived dataset resulted in a filtered list of 107 proteins (Figure 4C), including 52 shared across all groups, 16 unique to AA+MG132, and 16 common in both treatment conditions. In the FragPipe analysis, the filtered list comprises 208 proteins (Figure S3C), with 81 shared across all conditions, 21 specific to AA, 27 exclusive to AA+MG132, and 21 shared between both treated groups. Notably, both pipelines revealed multiple proteins with potential roles in proteasomal degradation. For example, previously discussed candidates such as UBR4, TRIP12, PSMA3 and PSMC2 were identified using this analysis strategy. FragPipe analysis also identified TRIM28 across all treatment conditions. Additionally, new proteins of interest were identified, including SQSTM1, DNAJA1, DNAJB1, HSPA1A and HSPA1B, which were identified in both MaxQuant and FragPipe. Although these proteins were present in all sample conditions, their levels were elevated in AA-treated samples, suggesting a possible role for AA in promoting these interactions. SQSTM1, also known as p62, is a scaffold protein that binds ubiquitinated proteins and primarily delivers them to the autophagosomes, or alternatively directs them to the proteasome, or sequesters them into aggresome-like structures(38). Interestingly, SQSTM1 has been shown to mediate the sequestration of ubiquitinated caspase-8, sensitizing HNSCC cells to ionizing radiation-mediated apoptosis(39). A similar mechanism may be triggered by AA treatment, facilitating delivery of ubiquitinated E7 for proteasomal degradation. Heat shock protein 40 (HSP40), including DNAJA1 and DNAJB1, are co-chaperones that cooperate with heat shock protein 70 (HSP70) family members, such as HSPA1A and HSPA1B, to promote proper protein folding(40,41). Notably, another member of this DNAJ family member, DNAJA3, has been reported to directly interact with E7(42). Furthermore, HSP70 proteins play a central role in proteostasis regulation(43). These findings raise the possibility that AA induces a conformational change in E7, enhancing E7 recognition by DNAJA1 or DNAJB1 to bind directly to E7, which may in turn recruit HSPA1A or HSPA1B to facilitate E7 degradation.

Together, our findings demonstrate the complementary strengths of MaxQuant and FragPipe pipelines, culminating in the overlapping identification of several unique candidate E7 interactors. This reinforces the robustness of our analytical approach and offers a solid foundation for future experimental validation. Although promising candidates have been identified, functional assays will be critical to determine their relevance in E7 degradation. To prioritize targets for further investigation, we next integrated complementary approaches to refine our candidate list.

### 3.4 Comparative analyses pinpoint HSP40 and HSP70 as potential mediators of E7 degradation via protein misfolding

Given the variety of available mass spectrometry (MS) data analysis strategies, we compared distinct stages of our analytical pipeline to identify key E7-interacting proteins. Network 1 included initial preprocessing, such as contaminant removal and filtering for valid values. Network 2 was generated using imputed, ANOVA-significant, z-scored data. For the final dataset, we excluded proteins with a NOK16/pWPI ratio below 1.05. Comparative analysis of these three networks revealed 49 (MaxQuant) and 77 (FragPipe) proteins that were shared across all three (Figure 5A). Additionally, most proteins were identified in at least two networks. This comparative analysis identified several proteins associated directly or indirectly with proteasomal degradation, including the E3 ubiquitin ligases TRIM28, RBBP6 and NOSIP, the HSPs DNAJA1, DNAJB1, HSP1A and HSP1B, and the proteasomal subunits PSMA6, PSMC1 and PSMD2. The overlapping results from both MaxQuant and FragPipe pipelines underscore the reliability of our approach in identifying high-confidence E7 interactors.

**Figure 5.**
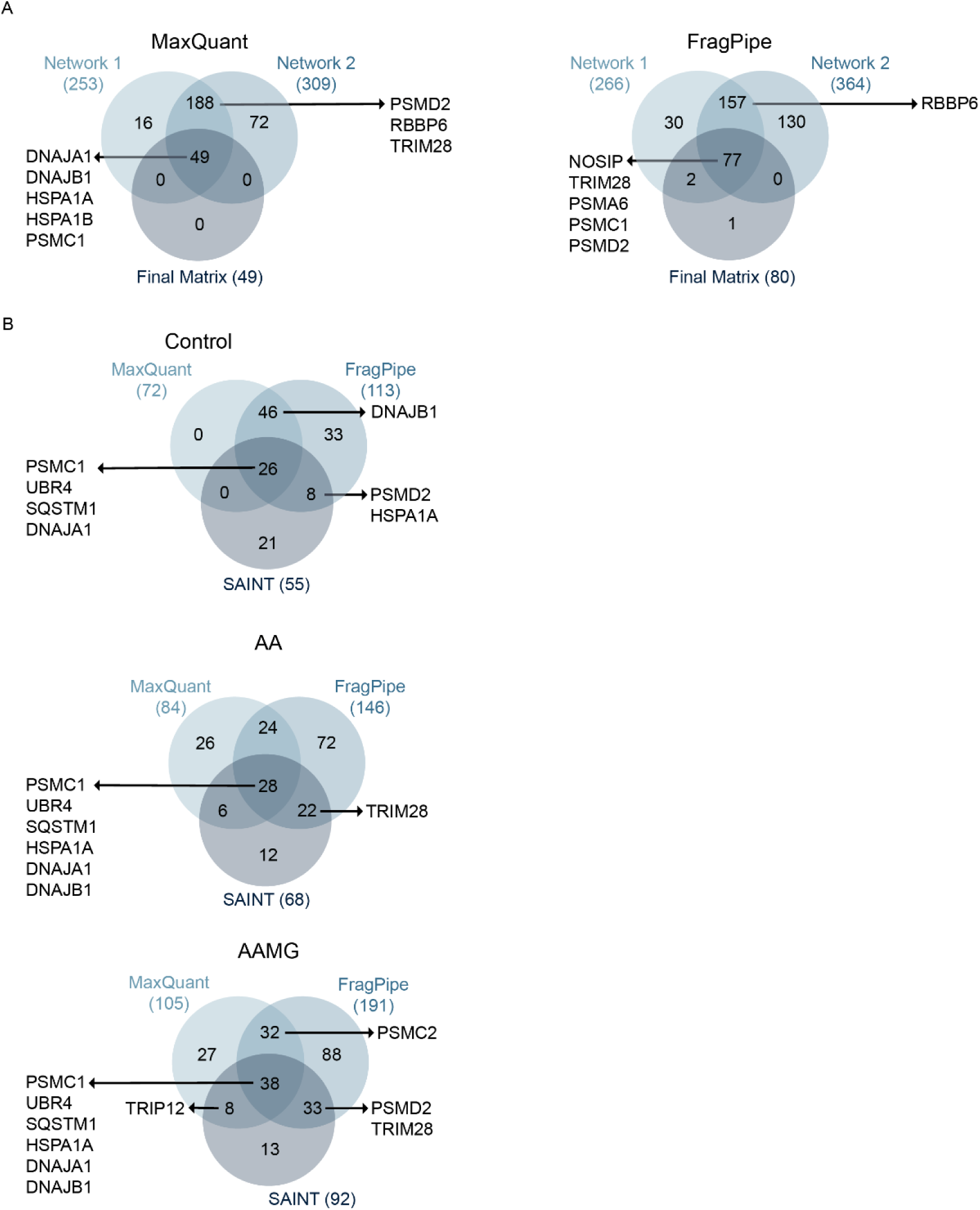
Multiple analytical pipelines consistently identify distinct E3 ligases and proteasomal subunits. a) Venn diagrams illustrate the overlap between protein networks and data matrices at different stages of MaxQuant (top) and FragPipe (bottom) analysis. Network 1 represents proteins obtained after initial preprocessing. Network 2 was generated from imputed ANOVA-significant z-scored data. The final matrix consists of proteins with a NOK16/pWPI ratio greater than 1.05. b) Venn diagrams comparing proteins identified by MaxQuant, FragPipe, and SAINT analyses across different conditions: Control (top), AA (middle) and AA+MG132 (bottom). Final matrices were used for the MaxQuant and FragPipe datasets.

To further enhance analytical depth, we used Significance Analysis of INTeractome (SAINT), a statistical method designed to differentiate true interactors from background proteins in interactome datasets(22). Data was filtered using thresholds of SAINT score (SP) > 2, fold change abundance (FCA) > 2, and a minimum abundance of 1.5. The resulting list was cross-referenced with our previous analyses (Figure 5B). This approach identified DNAJA1, DNAJB1, PSMC1, UBR4, SQSTM1, HSPA1A in all treatment conditions. Furthermore, TRIM28 was identified in both AA– and AAMG-treated samples, whereas TRIP12 and PSMC2 were only found in AAMG-treated samples. All these proteins have been consistently identified throughout our different analytical pipelines, highlighting their possible involvement in AA-mediated E7 degradation. Thus, our findings so far support a model in which AA binds directly to E7, potentially inducing a conformational change that increases its susceptibility to ubiquitination. This altered structure may enhance E7 recognition by E3 ligases or binding to chaperones, facilitating its ubiquitination and subsequent degradation via the proteasome.

HPV E7 is known to interact with different host proteins via different approaches. For instance, E7 binds to pRb using a LXCXE motif(44), to AP2 complex using the residues 25-YEQL-28(45), to CBP/p300 with high-affinity using the TAZ2 domain(46). Additionally, E7 is known to contain two CXXC zinc-binding domains in its C-terminus(47,48), which has been shown to allow nuclear import of HPV16 E7(49). The N-terminus of E7, on the other hand, is an intrinsically unstructured domain, which provides the protein flexibility and adaptability(50). Thus, to determine whether candidates could interact with E7 via similar motifs, we analyzed the sequences of each protein and assessed their relevance in a HPV infection environment. TRIM28 does not possess any of the domains and motifs mentioned before, but it does interact with MDM2 to promote p53 ubiquitination and degradation(51). While in an HPV-positive cancer cell p53 is targeted for degradation by the association of E6 and E6AP, reports have shown that in E7-expressing fibroblasts, MDM2 levels are increased(52), suggesting that indirectly modulate this alternative p53 degradation pathway. No evidence exists on whether MDM2 and E7 bind to each other, but our data raise the possibility that E7 may repress TRIM28-MDM2 complex in HPV-positive cells, favoring the E6-driven mechanism. Thus, in the presence of AA, the conformational changes in E7 could promote a E7-TRIM28-MDM2 complex that leads to the ubiquitination of E7 and subsequent degradation. Another E3 ligase of interest is UBR4. An interaction between E7 and UBR4 has been shown to induce PTPN14 degradation(53). In our recent study, UBR4 was also identified as a potential E6 interactor(12), and E6 and E7 were shown to interact directly to each other(54). This raises the possibility of UBR4 being able to mediate the degradation of both oncoproteins, contributing to the stabilization of pRb and p53, and activation of cell death induced by AA (11). TRIP12 is another candidate E3 ligase that was only identified in AAMG-treated samples. TRIP12 is known to interact with USP7, a deubiquitinase involved in DNA damage response and p53 stabilization. The TRIP12–USP7 axis is known to regulate each other’s expression(55). In the context of HPV, USP7 has been shown to stabilize E7, contributing to HPV-driven oncogenesis(56). Our data raise the possibility that AA could potentially destabilize the USP7-E7 interaction in favor of a TRIP12-E7, inducing E7 ubiquitination and degradation. Once ubiquitinated by a candidate E3 ligase, E7 could be delivered to the proteasome by the scaffold protein SQSTM1. SQSTM1 contains a long intrinsically disordered region, which serves as a binding site for several different proteins(57,58). Given this structural flexibility and its role in directing ubiquitinated proteins toward degradation pathways, it is conceivable that, in the presence of AA, SQSTM1 recognizes the ubiquitinated E7 and mediates its delivery to the proteasome.

Another possibility arises from the presence of a natively unfolded N-terminus in E7 protein(50), where AA could be crucial in changing E7 structural conformation and redirecting it to a protein misfolding-associated pathway. Interestingly, we identified HSP40 (DNAJA1 and DNAJB1), a co-chaperone protein that binds misfolded proteins and transfers them to HSP70 (HSPA1A), which was also identified here. HSP70 could then promote proper folding of E7, recruit an E3 ligase to ubiquitinate E7, or directly for degradation(59). This hypothesis is a strong one, as we identified three different proteins involved in such a process. Furthermore, our previous study on AA-effects on HPV16 E6 revealed that E6 is most likely targeted for degradation by E3 ligases, inducing its degradation. Thus, it would be interesting to investigate in future studies which oncoprotein is degraded first upon AA-treatment.

Through our integrative analysis across multiple MS pipelines and sequence motif analysis, and contextual protein function, we have identified TRIM28, TRIP12, and UBR4 as potential mediators of AA-induced degradation of HPV16 E7. Furthermore, an alternative degradation route may occur due to E7 interactions with members of the HSP40 and HSP70 families. These proteins act on distinct degradation pathways, including E3 ligase-mediated ubiquitination, scaffold-directed proteasomal targeting, and chaperone-assisted recognition of misfolded proteins. Thus, future studies are necessary to validate these interactions and clarify the degradation pathway of E7.

## 4 Conclusions

In this study, we used mass spectrometry-based proteomics to investigate the mechanism by which anisomelic acid promotes the degradation of the HPV16 E7 oncoprotein. Consistent with previous observations, we confirmed that AA treatment reduces E7 protein levels in HPV-transformed oral keratinocytes and that inhibition of proteasomal degradation prevents AA-mediated E7 degradation. Furthermore, the presence of AA leads to an increase in the ubiquitination of E7. Using CETSA, we provide compelling evidence that AA directly binds to E7, potentially inducing conformational changes that increase its susceptibility to ubiquitination. This finding highlights AA as a rare small molecule capable of binding an intrinsically disordered viral oncoprotein and shifting its structural stability toward degradation. Future studies focusing on characterizing the kinetics and binding affinity of the AA–E7 interaction will be beneficial to optimize the anisomelic acid therapeutic potential.

Through co-immunoprecipitation coupled with LC-MS/MS and integrative data analysis using MaxQuant, FragPipe, and SAINT, we identified several different high-confidence E7 interactors potentially involved in AA-mediated degradation. Among these, the E3 ligases UBR4, TRIP12 and TRIM28 emerged as the most promising candidates. Their consistent enrichment across multiple data pipelines, previous associations with HPV and/or roles in signaling pathways targeted by E7 strongly support their potential involvement in targeting E7 for degradation. Moreover, we identified the scaffold protein SQSTM1, known to mediate proteasomal delivery of ubiquitinated cargo, further suggesting a defined route of E7 clearance. Our proteomics data also revealed a potential alternative degradation mechanism involving protein misfolding. Specifically, we identified DNAJA1 and DNAJB1 (HSP40 co-chaperones) and HSPA1A (HSP70 chaperones) in AA-treated samples. These proteins typically recognize and process misfolded proteins, which are particularly relevant considering the intrinsically disordered nature of E7’s N-terminus. AA-induced conformational changes could expose E7 regions, leading to their recognition by HSP40/HSP70 complexes and eventual proteasomal degradation. The identification of multiple proteasomal subunits, including PSMC1, PMSC2 and PSMD2 further substantiates the proposed degradation model and clarifies the involvement of the 26S proteasome in this process. Together, our findings support a model in which AA binds directly to E7, altering its structural conformation, and enhances its recognition by specific E3 ligases or the HSP40/HSP70 machinery. While this model relied on integrating multiple analysis pipelines, biochemical validation is essential to confirm which pathway and specific proteins are involved in E7 degradation.

In conclusion, this study uncovers a potential molecular mechanism by which AA promotes the targeted degradation of HPV16 E7. These findings build upon our previous findings on E6 and suggest that AA may function as a specific therapeutic strategy against HPV-associated cancers and diseases. Future research should focus on validating the identified E3 ligases and chaperones in cellular and *in vivo* models and assessing whether AA analogues can be optimized for potency, selectivity, and clinical translation.

## Significance

This study investigates anisomelic acid as a potential therapeutic agent against HPV-positive cancers by elucidating its impact on the HPV16 E7 oncoprotein. We demonstrate that AA interacts with E7, promoting its ubiquitination and proteasomal degradation. By combining affinity purification-mass spectrometry with different proteomic analytical pipelines, we identified several candidate proteins that may be involved in AA-mediated E7 degradation. These findings provide new insights into how AA may target HPV E7 for degradation and identify promising targets for further functional validation.

## CRediT authorship contribution statement

**Michael Santos Silva:** Writing – review & editing, Writing – original draft, Visualization, Methodology, Investigation, Formal analysis, Data curation, Conceptualization. **Leila S. Coelho-Rato:** Writing – review & editing, Writing – original draft, Visualization, Methodology, Investigation, Formal analysis, Data curation. **Navid Delshad:** Writing – review & editing, Investigation. **Tatiana Tarkhova:** Investigation. **Joakim Edman:** Investigation. **Preethy Paul:** Writing – review & editing, Supervision, Funding acquisition. **Annika Meinander:** Writing – review & editing, Supervision. **John E. Eriksson:** Writing – review & editing, Supervision, Funding acquisition, Conceptualization.

## Acknowledgments

This research was supported by Jane Ja Aatos Erko Foundation, the Cancer Foundation Finland, Ida Montinin Säätiö and The Maud Kuistila Memorial Foundation. The research was carried out in Åbo Akademi University, and the Turku Bioscience Centre and Biocenter Finland research facilities. This work was further supported by InFLAMES Flagship Programme of the Academy of Finland (decision numbers: 337531, 357911).

## Data availability

The mass spectrometry proteomics data have been deposited to the ProteomeXchange Consortium via the PRIDE(60) partner repository with the dataset identifier PXD069948.

---

**Figure S1.**
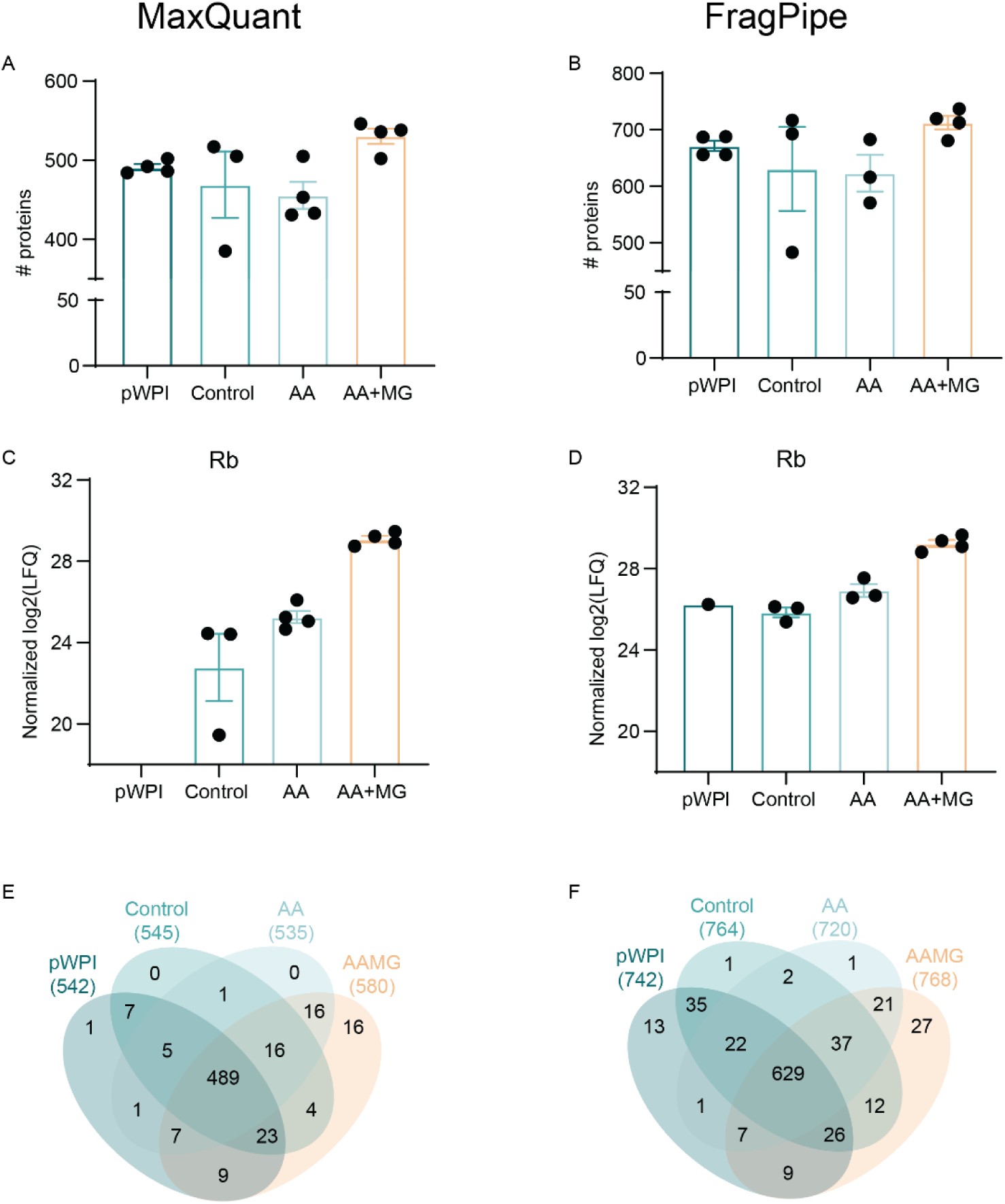
Analysis of unfiltered data in Perseus, using MaxQuant and FragPipe database search. a) and b) Bar charts showing the total number of proteins identified using MaxQuant (a) and FragPipe (b), prior to filtering steps. c) and d) Venn diagrams displaying the overlap of identified proteins across sample groups for MaxQuant (c) and FragPipe (d). e) and f) LFQ of pRb, a well-established E7 interactor, as detected in MaxQuant (e) and FragPipe (f).

**Figure S2.**
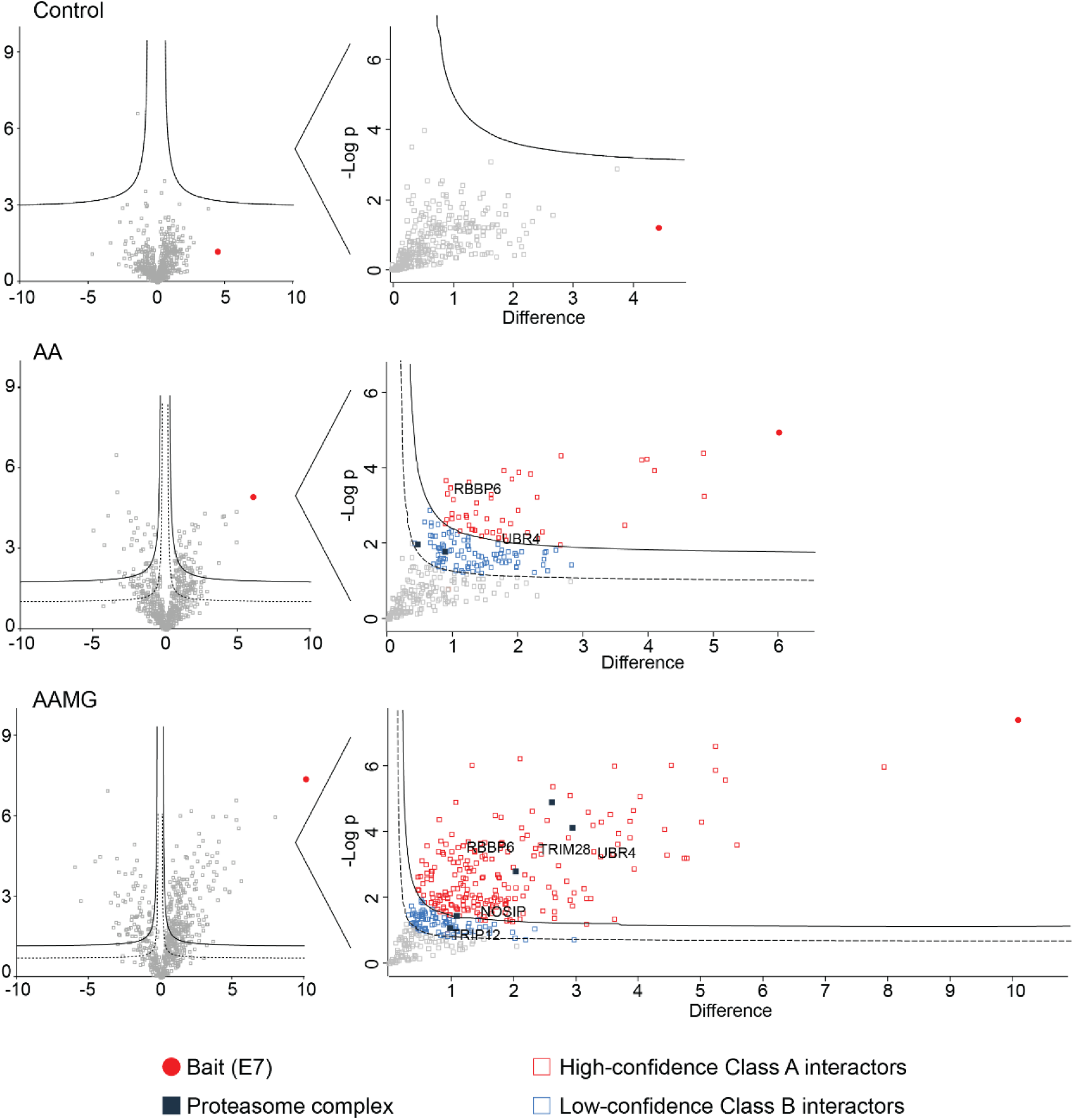
Hawaii plots identify proteasomal subunits and E3 ligases NOSIP, RBBP6, TRIM28, TRIP12, UBR4 as possible E7 interactors. a) Hawaii plots were generated for each condition: Control (top), AA (middle) and AA+MG132 (bottom). Interactors were classified based on permutation-based false discovery rate: high-confidence Class A (FDR < 0.01), positioned above the solid line and shown as red squares, and low-confidence Class B (FDR < 0.05), located between the solid and dashed lines and marked in light blue. The bait, E7, is highlighted in a filled red circle. Proteasome components are represented by dark blue squares.

**Figure S3.**
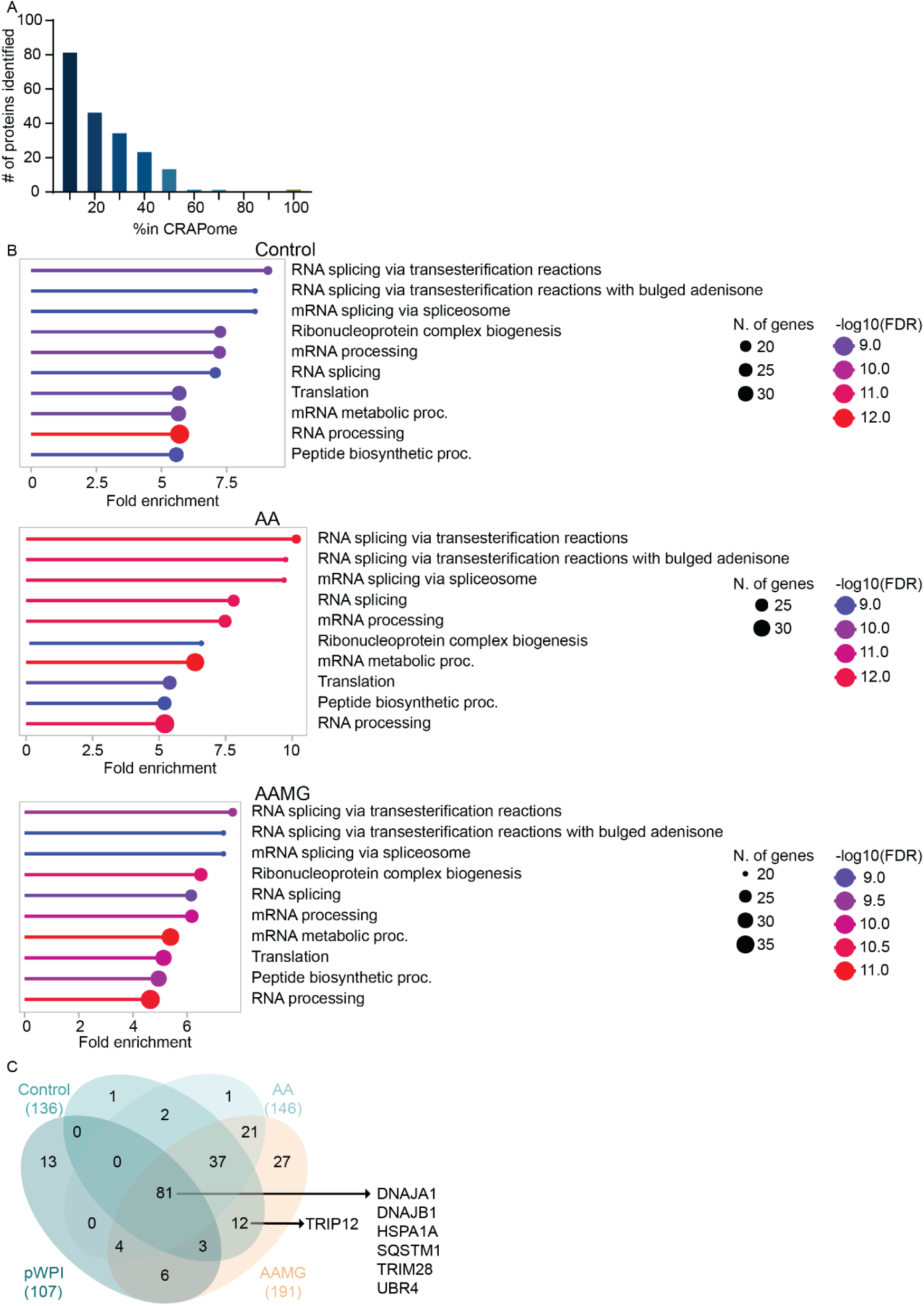
FragPipe and Perseus analyses reveal enrichment of E3 ligases and proteins involved in misfolded protein pathways in NOK16. a) The bar graph shows the number of proteins detected in the CRAPome database. A higher percentage indicates a more frequent occurrence of a protein. b) Dot plot illustrates enriched GO biological processes for each treatment condition. Circle size reflects the number of proteins associated with each GO term, while color represents –log₁₀(FDR). c) Venn diagram comparing protein identifications across experimental groups, following exclusion of proteins with a NOK16/pWPI ratio smaller than 1.05.

